# sRNARFTarget: A fast machine-learning-based approach for transcriptome-wide sRNA Target Prediction

**DOI:** 10.1101/2021.03.05.433963

**Authors:** Kratika Naskulwar, Lourdes Peña-Castillo

## Abstract

Bacterial small regulatory RNAs (sRNAs) are key regulators of gene expression in many processes related to adaptive responses. A multitude of sRNAs have been identified in many bacterial species; however, their function has yet to be elucidated. A key step to understand sRNAs function is to identify the mRNAs these sRNAs bind to. There are several computational methods for sRNA target prediction, and the most accurate one is CopraRNA which is based on comparative-genomics. However, species-specific sRNAs are quite common and CopraRNA cannot be used for these sRNAs. The most commonly used transcriptome-wide sRNA target prediction method and second-most-accurate method is IntaRNA. However, IntaRNA can take hours to run on a bacterial transcriptome. Here we present sRNARFTarget, a machine-learning-based method for transcriptome-wide sRNA target prediction applicable to any sRNA. We comparatively assessed the performance of sRNARFTarget, CopraRNA and IntaRNA in three bacterial species. Our results show that sRNARFTarget outperforms IntaRNA in terms of accuracy, ranking of true interacting pairs, and running time. However, CopraRNA substantially outperforms the other two programs in terms of accuracy. Thus, we suggest using CopraRNA when homolog sequences of the sRNA are available, and sRNARFTarget for transcriptome-wide prediction or for species-specific sRNAs. sRNARFTarget is available at https://github.com/BioinformaticsLabAtMUN/sRNARFTarget.

## 1 Introduction

sRNAs are bacterial small regulatory RNAs, usually less than 200 nucleotides in length, involved in several biological functions such as virulence, metabolism, and environmental stress response [16]. It is generally accepted that most bacteria have hundreds of sRNAs that regulate mRNA expression [20]. Many sRNAs exert their functions when they interact with mRNAs, and these interacting mRNAs are called the targets of the sRNAs. To understand the function and the regulatory networks of sRNAs, we first need to identify their targets.

There are several bioinformatics methods for sRNA target prediction such as CopraRNA [53], SPOT [22], TargetRNA2 [21], sTarPicker [56], and IntaRNA [43, 31]. CopraRNA, the most accurate method, requires sequence conservation of both sRNA and mRNA in at least four bacterial species, and must be run one sRNA at a time. The sequence conservation requirement makes CopraRNA unsuitable for species-specific sRNAs. Out of the programs that are not comparative genomic-based, IntaRNA and sTarPicker have been shown to achieve the best results in terms of the area under the ROC curve (AU-ROC) [36, 56]. IntaRNA is also the underlying algorithm of CopraRNA [53]. However, performing a transcriptome-wide sRNA target prediction on a bacterial transcriptome using IntaRNA might take several hours depending on the number of sRNAs and mRNAs investigated. Here we present sRNARF-Target, the first ML-based method that predicts the probability of interaction between an sRNA-mRNA pair. sRNARFTarget is generated using a random forest [7] trained on the trinucleotide frequency difference of sRNA-mRNA pairs. As sRNARFTarget bases its predictions on sequence alone, it can be applied to any sRNA-mRNA pair (i.e., does not require sequence conservation of either sRNA or mRNA). To train sRNARFTarget we collected known sRNA-mRNA interactions including those identified using RNA sequencing (RNA-seq) [49] approaches such as MAPS [23], GRIL-seq [19], CLASH [51] and RIL-seq [32]. To the best of our knowledge, this generated the largest data set of known sRNA-mRNA interactions from multiple bacteria available so far.

We comparatively assessed the performance of sRNARFTarget, CopraRNA and IntaRNA in terms of AUROC, ranking of confirmed interacting pairs, and running time using data from three bacterial species (*Escherichia coli, Pasteurella multocida* and *Synechocystis* sp PCC 6803). Our results show that CopraRNA is the most accurate and sRNARFTarget is the fastest of the three programs. Specifically, sRNARFTarget is on average 100 times faster than IntaRNA with the same or higher accuracy.

## 2 MATERIALS AND METHODS

### 2.1 Data collection

By searching in NCBI Pubmed, we identified studies listing confirmed sRNA-mRNA interactions including those identified by RNA-seq based methods (Table 2.1). We collected all sRNA-mRNA pairs listed in these studies and gathered roughly 2,400 sRNA-mRNA pairs from multiple bacteria.

**Table 1:** Training and benchmarking data characteristics

| Data | No. of species | No. of sRNAs | No. of pairs |
| --- | --- | --- | --- |
| Training | 37 | 176 | 745 |
| Benchmarking | 3 | 25 | 144 |

**Table 2:** Parameter per ML method used for grid-search CV.

| Method | Parameter | Values |
| --- | --- | --- |
| RF | n_estimators | [500, 600, 800, 1000] |
|  | max_features | ['sqrt', 'log2'] |
|  | max_depth | range(1, 11) |
| GB | n_estimators | [400, 500, 700, 1000] |
|  | max_features | ["log2", "sqrt"] |
|  | max_depth | range(1, 11) |
| KNN | n_neighbors | range(1, 50) |
|  | weights | ['distance', 'uniform'] |

The sRNA-mRNA pairs listed in the literature are in a variety of formats providing either sRNA mRNA names, sRNA - mRNA sequences, or sRNA mRNA genomic locations. We used the sequences directly if they were provided (e.g., sTarBase3.0 [50]). For other datasets, we created a file containing Entrez genome accession number, sRNA name and target mRNA name per sRNA-mRNA pair (Supplementary Tables 2 and table 3).

| Bacterium | Data source |
| --- | --- |
| *Escherichia coli* | [24, 33, 34, 36, 48, 51] |
| *Pseudomonas aeruginosa* | [10, 19, 40] |
| *Burkholderia cepacia* | [40] |
| *Pasteurella multocida* | [17] |
| *Salmonella* | [13, 44] |
| *Mycobacterium tuberculosis* | [30] |
| *Synechocystis* | [14, 15] |
| Multiple bacteria | [6, 50] |

Our first data preprocessing step was to remove any duplicate pairs. We then divided the collected data into training data and validation data. 102, 22, and 20 sRNA-mRNA pairs from *Escherichia coli* [36], *Pasteurella multocida* [17] and *Synechocystis* [15] [14] respectively, were held-out for benchmarking. The remaining data was used for training the models (Table 1, and Supplementary Tables 1-3).

To get the complete sRNA and mRNA sequences, we wrote two Nextflow (version 0.32.0) [9] pipelines. The first pipeline finds whether the sRNAs and mRNAs names exist in the NCBI Gene database using the esearch function of Entrez direct [46] and generates a table containing sRNA-mRNA pairs found in the NCBI Gene database. Then our second pipeline gets the sRNA/mRNA sequences using esearch from Entrez direct, and bedtools (version 2.27.1) [41]. All Nextflow pipelines used are available at https://github.com/BioinformaticsLabAtMUN/sRNARFTarget.

### 2.2 Machine learning model generation

We generated models for sRNA target prediction using three ML methods, namely, Random Forest (RF) [7], K-nearest neighbours (KNN) [35] and gradient boosting (GB) [12] using scikit-learn [38] functions to implement these classifiers.

#### 2.2.1 Training Data

We used *k*-mer frequency difference, and secondary structure distances as features to train the machine learning models. To calculate *k*-mer frequency difference, one first has to separately compute *k*-mer frequency for both sequences (sRNA and mRNA), and then calculate for every *k*-mer *i, f*_*i,mRNA*_ − *f*_*i,sRNA*_ where *f*_*i,s*_ is the frequency of *k*-mer *i* in sequence *s*. To obtain *k*-mer frequency and then *k*-mer frequency difference we ran another Nextflow pipeline using scikit-bio (version 0.5.5) [4] in Python (version 3.7.4). We used *k* equal to 3 and 4, which corresponds to 64 and 256 *k*-mers, respectively. We obtained predicted secondary structures of sRNAs and mRNAs using CentroidFold (version 0.0.16) [18] with default values. Then we calculated seven distances between sRNA and mRNA secondary structures using RNAdistance (version 2.4.13) [27] program with default values indicating with the -D parameter the distance to calculate (F, H, W, C, h, w, or c).

After processing, our training data contained 745 interacting sRNA-mRNA pairs collected from the literature (Supplementary Table 2). We created negative instances by randomly swapping the sRNAs in the sRNA-mRNA pairs. Basically, negative instances are sRNA-mRNA pairs where there is no experimental evidence for interaction. The use of nonannotated sRNA-mRNA pairs as negative instances gives a conservative estimate of the performance of the models (some predictions considered false positives might in fact be true positives). In total, we had 1490 sRNA-mRNA pairs (745 positives and 745 negatives) for training the ML models.

In sum, we have a balanced training data with 1,490 instances for a binary classification task, and explore four feature sets with a) 64 (trinucleotide frequency difference), b) 71 (trinucleotide frequency difference plus seven distances), c) 256 (tetra-nucleotide frequency difference), and d) 261 (tetra-nucleotide frequency difference plus seven distances) attributes.

#### 2.2.2 Model Training

We used grid-search cross-validation (CV) of scikitlearn to get the best parameters per ML method. Table 2 shows the parameter ranges used in grid-search CV. We did 10-fold stratified CV to ensure balanced class distribution in each fold and used the area under the ROC curve (AUROC) to evaluate model performance. Additionally, we used R importance function [2] based on mean decrease in accuracy to get the feature importance, and filtered out any feature with a mean decrease in accuracy ≤ 0.

#### 2.2.3 Model Selection

We calculated sRNA-mRNA secondary structure distances to explore whether these features will increase AUROC and added them as features together with the trinucleotide frequency difference or the tetranucleotide frequency difference. Thus, we trained models using either trinucleotide frequency difference (64 features), tetra-nucleotide frequency difference (256 features), trinucleotide frequency difference plus seven distances, or tetra-nucleotide frequency difference plus seven distances. For each of the four sets of features, we found the optimal parameter setting per classifier using grid search CV and compared the models’ performance in terms of 10-fold CV AUROC. We selected the model with the highest AUROC, and saved this model to be used by the Nextflow pipeline implementing sRNARFTarget.

### 2.3 sRNARFTarget Nextflow pipeline

We wrote a Nextflow pipeline that uses our best model for sRNA target prediction. The pipeline takes sRNA and mRNA FASTA files as input, creates all possible sRNA-mRNA pairs, obtains the *k*-mer frequency for both sRNA and mRNA, and calculates the *k*-mer frequency difference by subtracting sRNA frequency from mRNA frequency using pandas (version 0.25.1) [37, 52] subtract function. Then the saved best model is loaded and predictions for all pairs are generated. The final result of the pipeline is a CSV file containing predicted probabilities of sRNA-mRNA interaction sorted in descending order with the sRNA-mRNA ID (see Supplementary Figure 1 for workflow of sRNARFTarget program). Additionally, a file containing the features for all sRNA and mRNA pairs is also created. This file is used by the interpretability programs.

### 2.4 sRNARFTarget Interpretability

We wrote two Python scripts using SHAP (version 0.35.0) [29] and pyCeterisParibus (version 0.5.2) [5] packages to facilitate the interpretation of predictions generated by sRNARFTarget (Supplementary Figure 2). Both scripts use the feature file generated by sRNARFTarget to get the features for the pair of interest. sRNARFTarget SHAP uses TreeExplainer of SHAP to create an explainer. Then it calculates the SHAP values for a given observation and generates SHAP’s decision, waterfall and force plots for interpretation. sRNARFTarget CP creates the explainer using training data and calculates ceteris paribus profiles for a chosen feature for given sRNA-mRNA pair. It then generates a plot of the calculated profiles for the selected feature.

### 2.5 Benchmarking

Previous comparative assessments of sRNA target prediction programs [56, 36, 22] reported four programs (CopraRNA, IntaRNA, SPOT and sTarPicker) as the most accurate programs, with CopraRNA been the most accurate program. SPOT is reported to be comparable to CopraRNA; however, we were unable to run SPOT locally and running SPOT through Amazon Web Services (AWS) requires payment [3]. Additionally, sTarPicker program is no longer available. Therefore, we included CopraRNA and IntaRNA in our benchmark.

The data used for independent benchmarking have 22 sRNAs and 102 confirmed interacting sRNA-mRNA pairs for *E. coli* [36], one sRNA and 22 confirmed sRNA-mRNA pairs for *P. multocida* [17], and two sRNAs and 20 pairs for *Synechocystis* bacteria [15, 14]. These data were not used for training. We extracted the sequences for 22 sRNAs of *E. coli* using our Nextflow pipeline as described above. For all other sRNAs, we fetched the sequence directly from the NCBI nucleotide database based on the locations provided in the corresponding manuscript. The location of *isar1* sRNA was taken as reported in [15]. The location of *psrR1* sRNA (1671919-1672052) was confirmed by oral communication with the author of [14]. Finally, *gcvB* sRNA location was obtained from [17]. As we wanted to perform transcriptome-wide prediction of sRNA targets, we collected genomic location for all the mRNAs belonging to each bacterium directly from NCBI. We then obtained the sequences for these mRNAs using our Nextflow pipeline. In the case of CopraRNA, if predictions for a given *E. coli* sRNA were already available in CopraRNA web server, we used the available predictions. Otherwise, to find homologs for *E. coli* sRNAs, we used GLASSgo - sRNA Homolog Finder [28]. Additionally, we used the homologs provided in [15] and [14] for *isar1* and *psrR1* sRNAs of *Synechocystis*. For *gcvB* sRNA of *P. multocida*, we retrieved homolog sRNAs from NCBI.

We downloaded IntaRNA (version 3.1.0.2) source code from [1], installed it locally, and executed it with default values from the command line. To obtain a total execution time for IntaRNA, we created a Nextflow pipeline to run IntaRNA’s two steps: 1) getting the interaction energy and 2) calculating the p-values for the interaction energy. We ran sRNARFTarget and IntaRNA from the Linux command line (system specifications are: one processor, processor speed 2.2 GHz, 4 cores and 16 GB RAM). CopraRNA (version 2.1.2) was run from its web server (http://rna.informatik.uni-freiburg.de/CopraRNA/Input.jsp,version4.8.2).

After running the programs, we standardized their results by assigning corresponding classes to all predictions (1 to confirmed interacting sRNA-mRNA pairs and 0 to all other sRNA-mRNA pairs) and using predicted interaction probability for all programs. CopraRNA and IntaRNA output p-values where lower p-values indicate higher predicted likelihood of interaction. Thus, we subtracted CopraRNA and IntaRNA p-values from 1 to obtain predicted interaction probability. Additionally, for all three programs we rounded the predicted interaction probability to 5 decimals. To eliminate the duplicate entries from CopraRNA result, we wrote an R (version 3.5.1) script to get the most significant p-value (lowest p-value) for each sRNA-mRNA pair, and remove all other entries. As CopraRNA did not produce a prediction for all possible sRNA-mRNA pairs. We wrote an R script to get the common pairs predicted by the three programs so that all three programs were evaluated on the same mRNA-sRNA pairs (Table 3).

**Table 3:** Final benchmarking dataset used for all three programs. The table lists the genome accession used, the number of sRNAs, the number of mRNAs, the number of confirmed interacting pairs (P), and the number of pairs considered non-interacting (N) per bacterial species (from top to bottom: *E. coli, Synechocystis* and *P. multocida*).

| Accession | sRNAs | mRNAs | P | N |
| --- | --- | --- | --- | --- |
| NC_000913.3 | 22 | 4240 | 101 | 92348 |
| NC_000911.1 | 2 | 3179 | 20 | 6324 |
| NC_002663.1 | 1 | 1804 | 22 | 1781 |

## 3 RESULTS AND DISCUSSION

### 3.1 Selection of sRNARFTarget ML model

We adopted the idea of using sequence-derived features such as *k*-mer frequency from previous studies [55, 25, 39, 8]. As sRNAs bind mRNAs through base pairing [47], we hypothesized that *k*-mer frequency difference might capture base pairing potential between mRNA and sRNA for the classifiers to use. Thus, we created feature sets using trinucleotide and tetra-nucleotide frequency difference. We started with trinucleotide composition, and as the performance decreased with tetra-nucleotide composition, we decided not to go beyond tetra-nucleotide composition.

Table 4 shows the performance in terms of AU-ROC of the best model per classifier when trained using trinucleotide frequency difference and tetranucleotide frequency difference. AUROC achieved with trinucleotide frequency difference was higher than the AUROC achieved with tetra-nucleotide frequency difference. With trinucleotide frequency difference, the model with the best performance in terms of AUROC was the RF one with 0.67, followed by GB with 0.66, and then KNN with 0.63.

RNA secondary structures are associated with the regulation of mRNA [11]. Previous studies [45, 8] used secondary structure information for prediction of sRNA-mRNA interaction and non-coding RNAs. As the secondary structure of both sRNA and mRNA affects their binding [54], we decided to assess whether including secondary structure distances as features together with the tri(tetra)-nucleotide frequency difference improved performance in terms of AUROC. However, including predicted secondary structure distances to the feature set did not increase the models’ performance. When including secondary structure distances as features in addition to trinucleotide frequency difference, the AUROC was unchanged for RF (AUROC 0.67), dropped by more than half for KNN (AUROC 0.27) and went slightly up for gradient boosting (AUROC 0.67). Similarly, adding secondary structure distance features with tetra-nucleotide frequency difference features had little to no effect on model performance (Supplementary Table 4). As adding distance features did not substantially improve models’ performance but dramatically increased the time required to extract the features from seconds to hours (due to the prediction of RNA secondary structure using CentroidFold), we decided against using the distance features in our final model.

**Table 4:** 10-fold CV AUROC for the best model per classifier trained on sequence-derived features (trinucleotide frequency difference and tetra-nucleotide frequency difference) of 1490 sRNA-mRNA pairs.

| Models | AUROC<br>(mean ± standard deviation) | |
| --- | --- | --- |
|  | Tri nt.<br>difference | Tetra nt.<br>difference |
| **RF** | 0.67 ± 0.03 | 0.31 ± 0 |
| **GB** | 0.66 ± 0.03 | 0.32 ± 0 |
| **KNN** | 0.63 ± 0.03 | 0.45 ± 0.01 |

RF and GB models were comparable in terms of AUROC; however, the RF model was much faster to train than GB. Thus, we decided to train our final model on the 1490 sRNA-mRNA pairs using RF and included this model in the sRNARFTarget pipeline. The parameters to create this model are 500 trees (n estimators), log2 of features for split (max features), and a maximum depth of the trees of 9 (max depth). From now on, we will refer to this final RF model as sRNARFTarget. Supplementary Figure 3 shows the 10-fold CV ROC curve of sRNARFTarget and Supplementary Figure 4 shows its top 30 most important features.

### 3.2 Interpreting sRNARFTarget predictions

To facilitate the interpretation of sRNARFTarget predictions, we have implemented two pipelines (sRNARFTarget SHAP and sRNARFTarget CP) to apply interpretability programs to sRNARFTarget predictions. To illustrate the functionality of these pipelines, we discuss interpretability plots generated for *isaR1-petF* confirmed interacting pair of *Synechocystis*. SHAP’s decision plot shows how the model reached its decision (Supplementary Figure 5). The waterfall plot suggests that the value of feature GGC lowers the probability of interaction for this pair, and that this feature has the highest relevance for this observation (Supplementary Figure 6). Force plot shows that features ACC and AAT are pushing sRNARFTarget to output higher interaction probability for this pair (Supplementary Figure 7). To gain insight on how a different value for the feature GGC impacts the output of sRNARFTarget for this pair, we looked at the ceteris paribus plot for feature GGC for *isaR1-petF* pair from *Synechocystis* (Supplementary Figure 8). It shows sRNARFTarget’s prediction for different values of GGC when all other feature values remain constant. These plots can help pinpoint the sequence segments (trinucleotides) that contribute more to a specific sRNA-mRNA interaction.

### 3.3 Benchmark on independent data set

First, we assessed the performance of sRNARFTarget, CopraRNA and IntaRNA in terms of AUROC on data from three bacterial species: *E. coli* (gammaproteobacteria), *Synechocystis* (cyanobacteria) and, *P. multocida* (gammaproteobacteria). These data were not used for training. The *E. coli* 102 confirmed sRNA-mRNA pairs were the same used in the assessment performed by Pain et al [36]. We performed transcriptome-wide predictions; i.e., the methods have to infer interaction probability for all possible sRNA-mRNA pairs. Note that this is a conservative assessment as there might be true sRNA-mRNA interacting pairs that have not been confirmed yet and are considered false positives in the evaluation. Figs. 1, 2 and 3 show the ROC curve for *E. coli, Synechocystis* and *P. multocida*, respectively. Table 5 shows the AUROC for the three programs per bacterium. Across the three bacterial species, CopraRNA has the highest AUROC followed by sRNARFTarget and then IntaRNA. All programs show a decrease in AUROC on *P. multocida* data. As the data used is highly unbalanced (Table 3), we also obtained the Precision-Recall curves (PRC) (Supplementary Figures 9-11). As it can be seen from the PRC curves and the AUPRC shown in Supplementary Table 5,there is still room for improving the precision of computational transcriptome-wide sRNA target prediction. This result is similar to that obtained by Pain et al [36].

**Figure 1:**
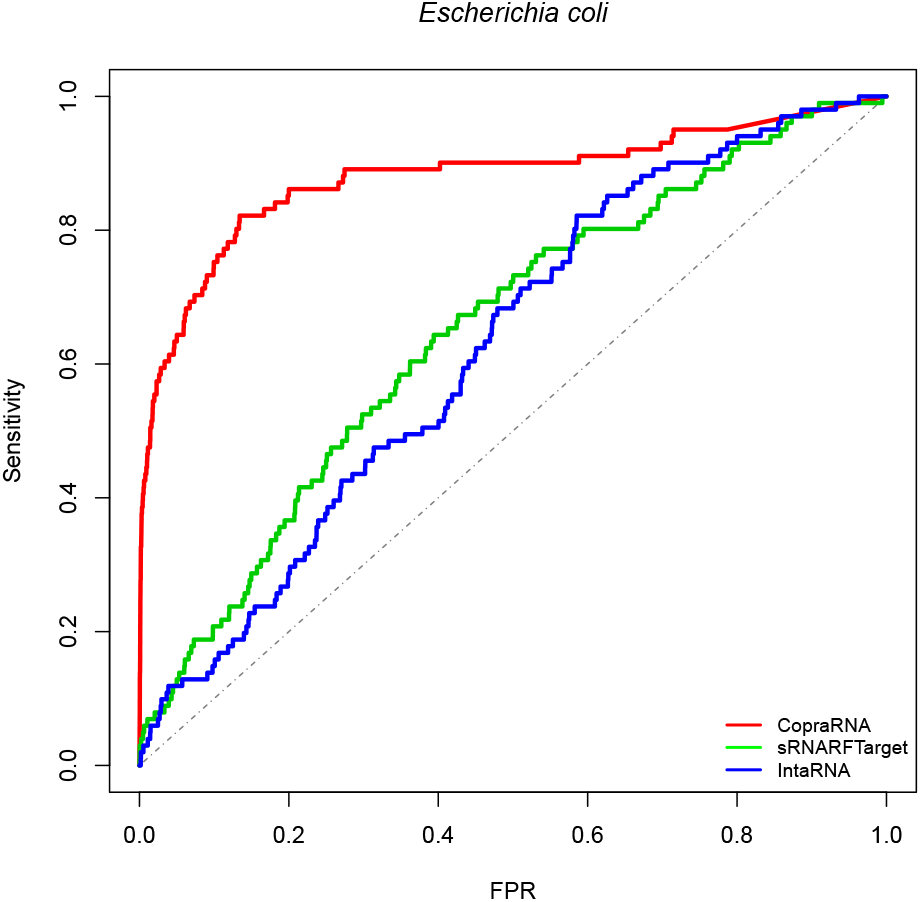
ROC curve for the three programs on *Escherichia coli* data. The plot shows the sensitivity (also called recall or true positive rate) as a function of the false positive rate (FPR). The dash line indicates random classifier performance.

**Figure 2:**
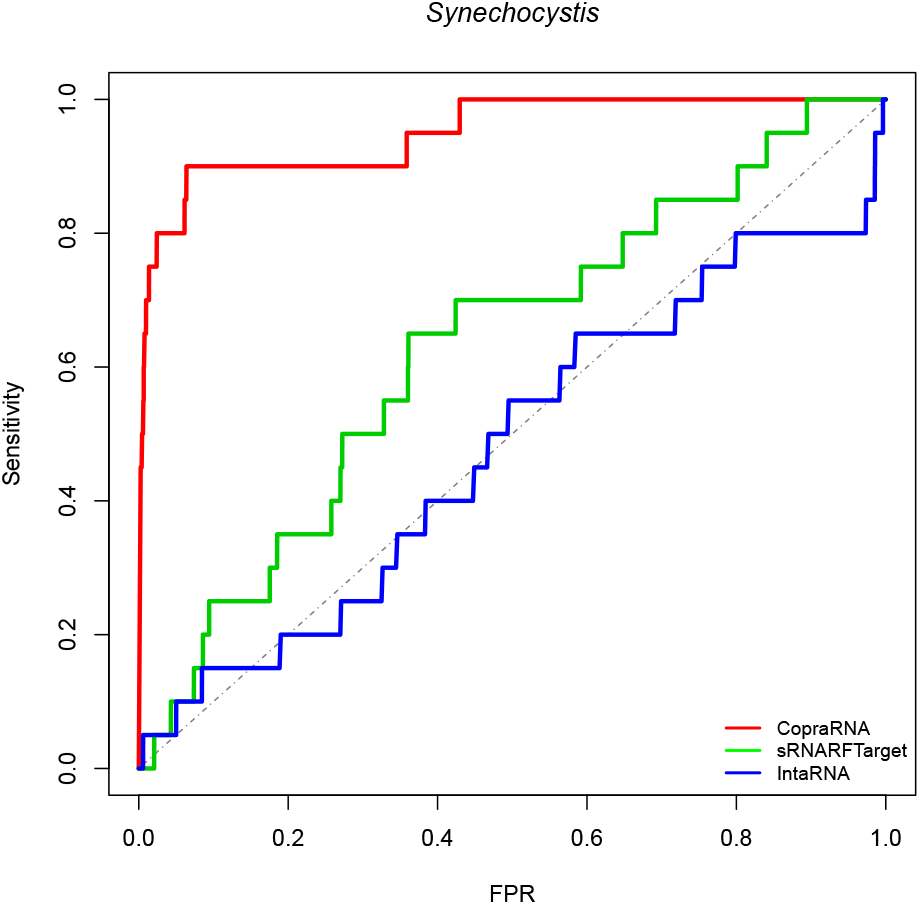
ROC curve for the three programs on *Synechocystis* data. The plot shows the sensitivity (also called recall or true positive rate) as a function of the false positive rate (FPR). The dash line indicates random classifier performance.

**Figure 3:**
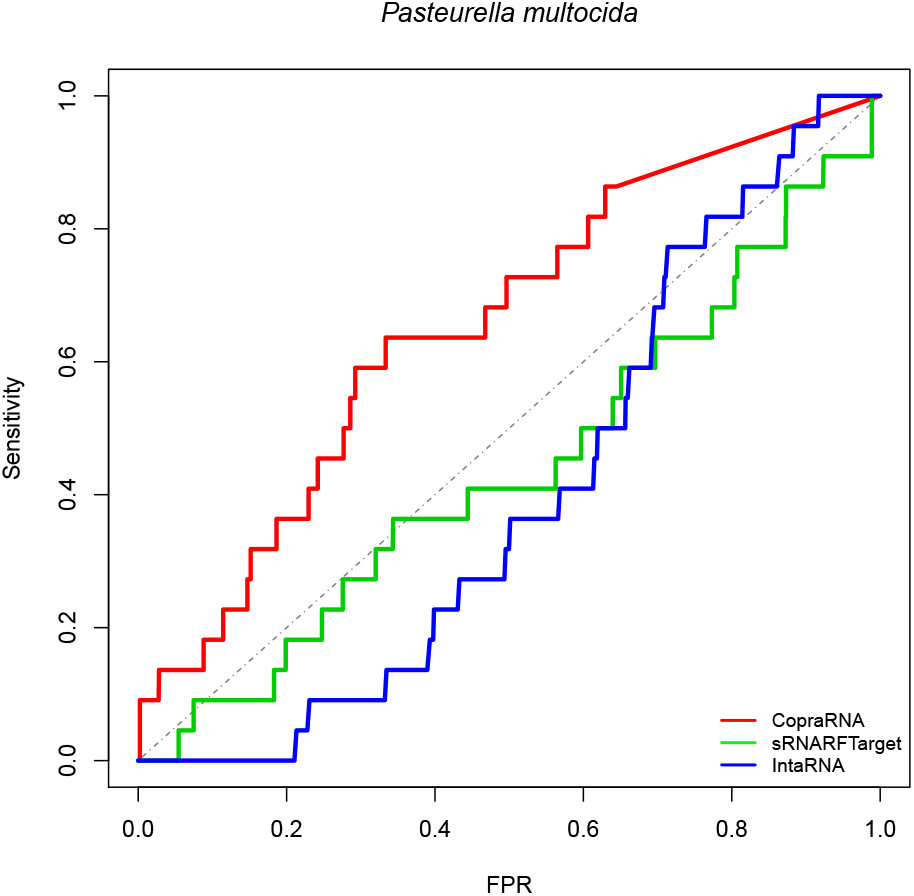
ROC curve for the three programs on *Pasteurella multocida* data. The plot shows the sensitivity (also called recall or true positive rate) as a function of the false positive rate (FPR). The dash line indicates random classifier performance.

**Table 5:** AUROC obtained on each bacterial species included in the benchmark for all three programs assessed.

| Bacterium | CopraRNA | sRNARFTarget | IntaRNA |
| --- | --- | --- | --- |
| *E. coli* | 0.88 | 0.65 | 0.62 |
| *Synechocystis* | 0.95 | 0.63 | 0.48 |
| *P. multocida* | 0.65 | 0.44 | 0.40 |
| **Average ± sd** | **0.83 ± 0.16** | **0.57 ± 0.12** | **0.50 ± 0.11** |

Next, we looked at the rank distribution of confirmed interacting pairs per bacterium. Ideally, actual interacting pairs should have lower rank than non-interacting pairs, as a lower rank indicates that the program predicts with higher confidence that a given sRNA-mRNA pair is an actual interacting pair. To visualize program performance in terms of ranking of confirmed interacting pairs, we generated violin plots showing the rank distribution of confirmed interacting sRNA-mRNA pairs. The shape surrounding the box plots indicates the data density for different rank values. The horizontal bar inside the box shows the median rank of the confirmed interacting pairs. Fig. 4 shows the violin box plot for *E. coli*. CopraRNA has a lower median rank followed by sRNARFTarget and then IntaRNA. The shape of CopraRNA suggests that most of the confirmed interacting pairs are ranked before all other pairs. The shape of the plot for sRNARFTarget suggests that it has more confirmed interacting pairs with lower ranks than IntaRNA. We compared the rank distributions using the Mann-Whitney test (Fig. 4). The p-values obtained indicate that CopraRNA’s median rank of interacting pairs is significantly lower than sRNARFTarget’s median rank, and that sRNARF-Target’s median rank is significantly lower than IntaRNA’s median rank.

**Figure 4:**
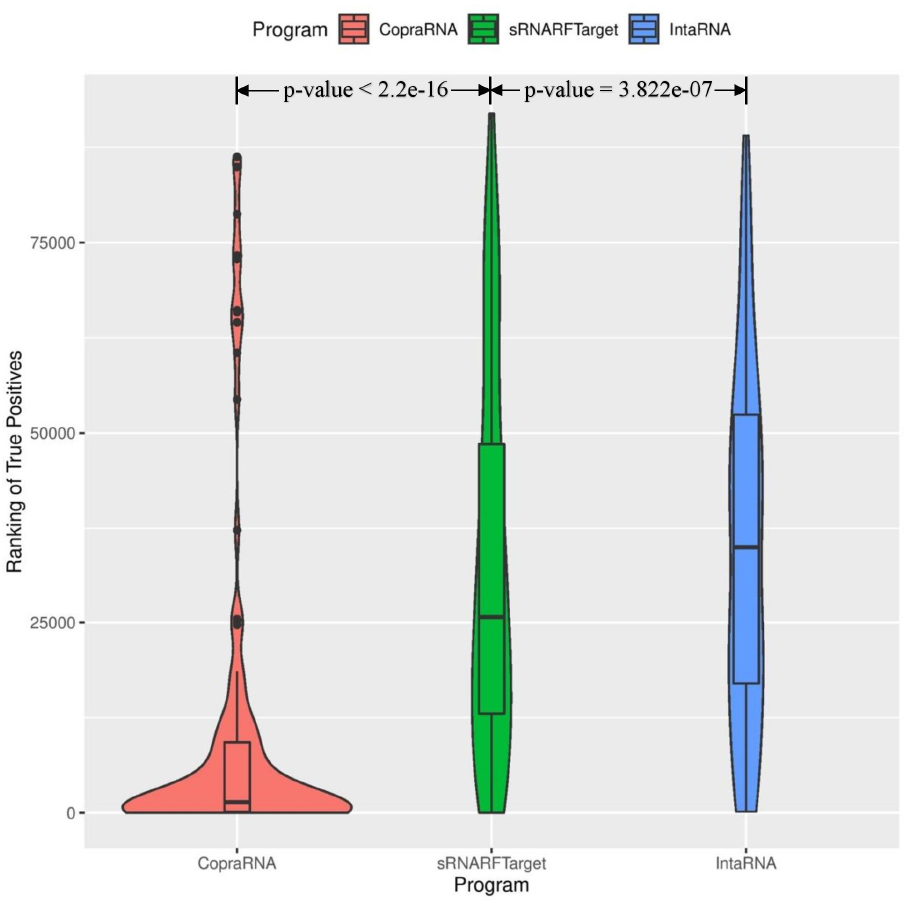
Rank (lower = better) distribution of 102 *Escherichia coli* confirmed interacting pairs. The violin plot for each program shows the data density for different rank values and the horizontal line inside each box indicates the median rank of confirmed interacting pairs.

Figs. 5 and 6 show the violin plots for *Synechocystis* and *P. multocida*, respectively. For these two bacterial species as well, the median rank of confirmed interacting pairs is the lowest in CopraRNA’s predictions, followed by sRNARFTarget and then IntaRNA. All three programs found it more difficult to distinguish true interacting pairs in *P. multocida* and ranked confirmed interacting pairs with higher ranks (Fig. 6) than for the other two bacteria. Nevertheless CopraRNA still ranks confirmed interacting pairs significantly lower than sRNARFTarget (p-value = 2.15e-05), and sRNARFTarget ranks true interacting pairs lower than IntaRNA (p-value = 0.056).

**Figure 5:**
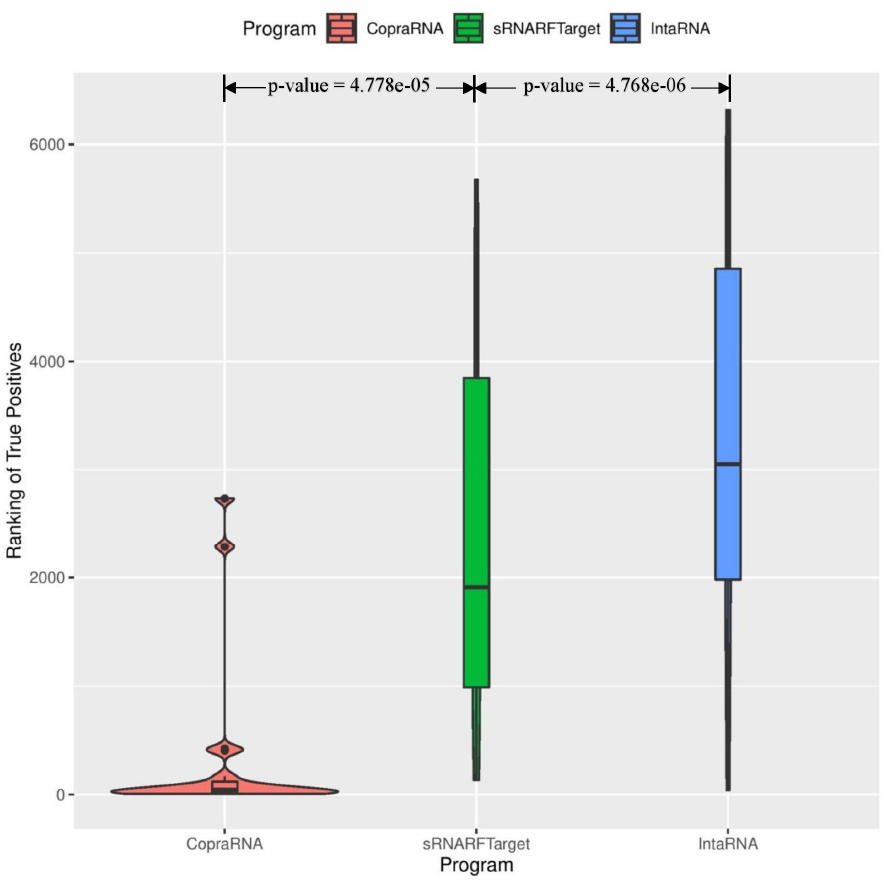
Rank (lower = better) distribution of 22 *Synechocystis* confirmed interacting pairs. The violin plot for each program shows the data density for different rank values and the horizontal line inside each box indicates the median rank of confirmed interacting pairs.

**Figure 6:**
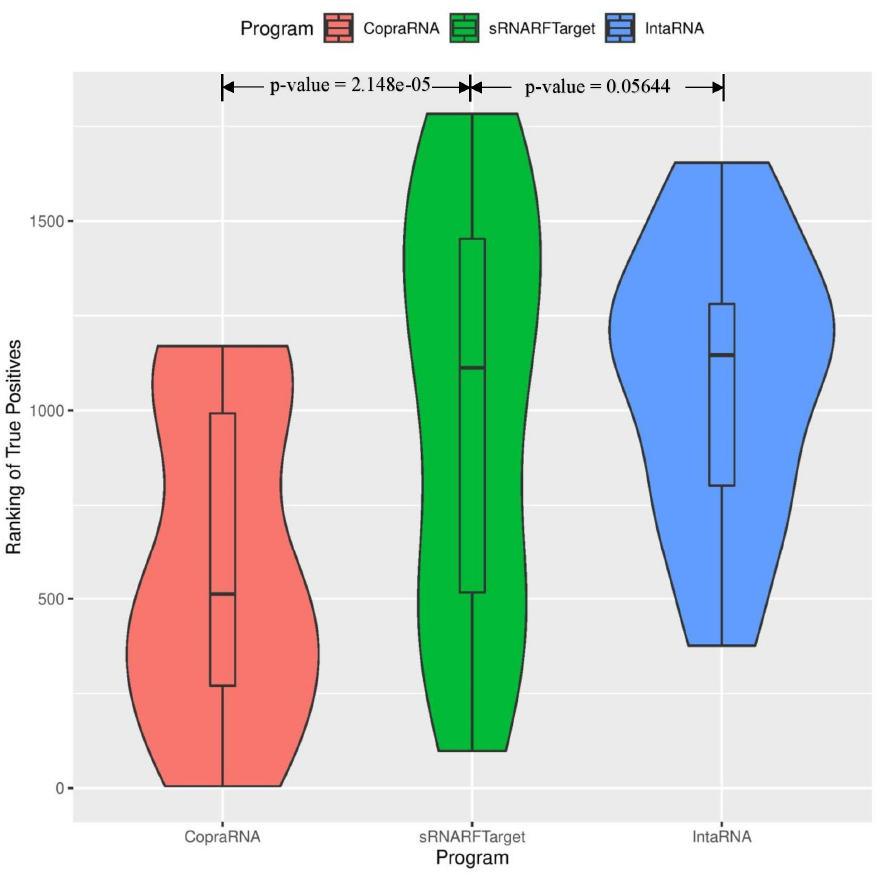
Rank (lower = better) distribution of 20 *Pasteurella multocida* confirmed interacting pairs. The violin plot for each program shows the data density for different rank values and the horizontal line inside each box indicates the median rank of confirmed interacting pairs.

Lastly, we plotted the percentage of confirmed interacting sRNA-mRNA pairs predicted among a certain percentage of top predicted interacting pairs. To create these plots, we took the top 10% predictions for each program, counted the number of confirmed interacting pairs among these predictions, and calculated the percentage of true positives (recall) among the top 10% predictions. Then iteratively increased the percentage of top predictions by 10% and repeated the process described above until all predictions (100%) were taken into account. We plotted the percentage of predictions on the x-axis and the percentage of confirmed interacting pairs (recall) on the y-axis. Fig. 7 shows this plot for *E. coli*. In the top 10% predictions, CopraRNA predicted 74% of confirmed interacting pairs, sRNARF-Target predicted 21% of these pairs, and IntaRNA predicted 14%. Among the top 50% predicted interacting pairs on *Synechocystis*, CopraRNA predicted 100% of the confirmed interacting pairs, sRNARF-Target predicted 70% of these pairs and IntaRNA predicted 55% (Fig. 8). In the top 20% predictions for *P. multocida*, CopraRNA predicted 18% of confirmed interacting pairs, sRNARFTarget was able to predict 10% of these pairs, and IntaRNA did not predict any confirmed interacting pair (Fig. 9). Thus, sRNARFTarget recovers more verified sRNA-mRNA interacting pairs than IntaRNA.

**Figure 7:**
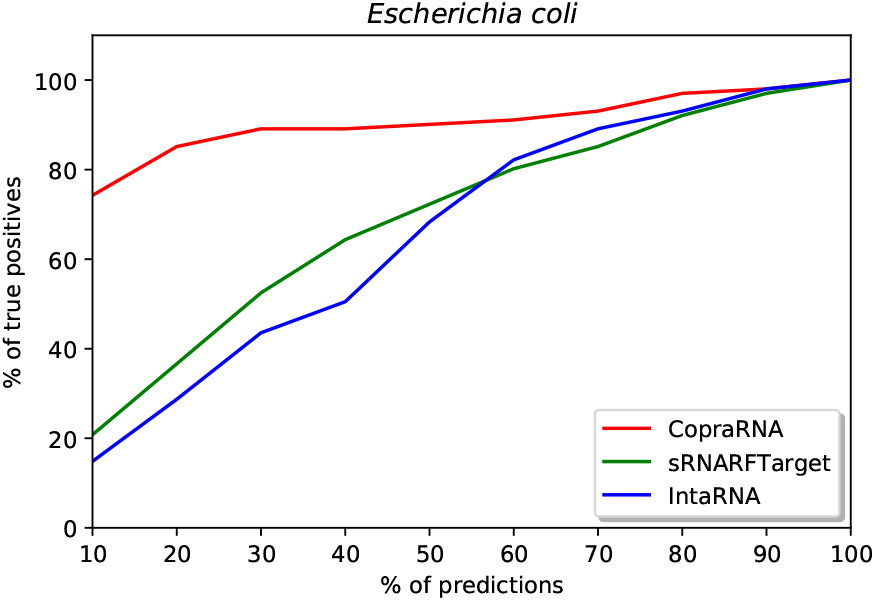
Percentage of *Escherichia coli* confirmed interacting sRNA-mRNA pairs (recall) as a function of percentage top predicted interacting pairs.

**Figure 8:**
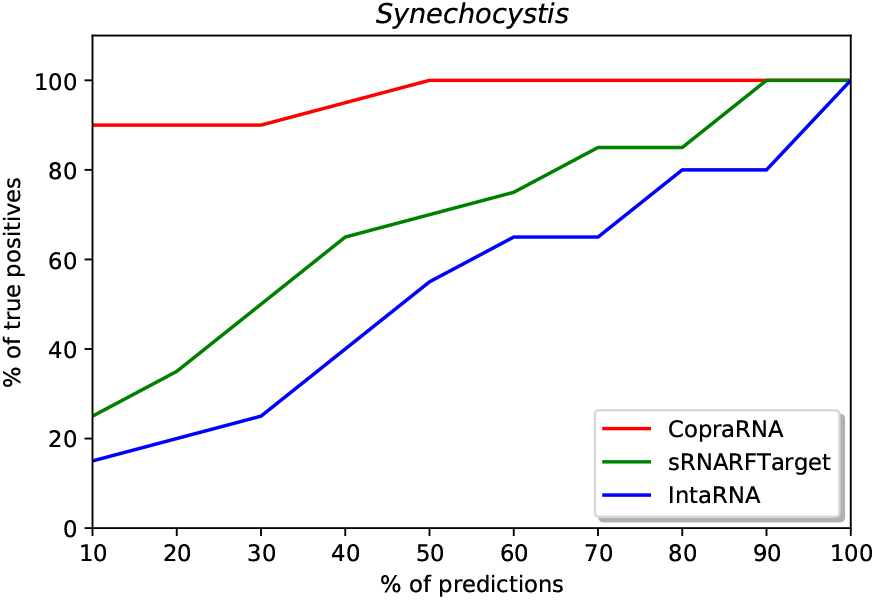
Percentage of *Synechocystis* confirmed interacting sRNA-mRNA pairs (recall) as a function of percentage top predicted interacting pairs.

**Figure 9:**
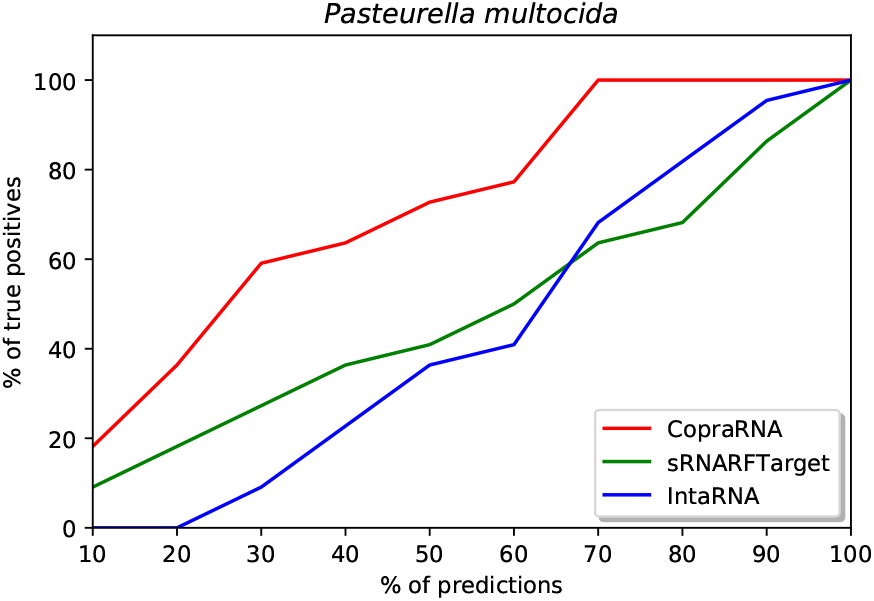
Percentage of *Pasteurella multocida* confirmed interacting sRNA-mRNA pairs (recall) as a function of percentage top predicted interacting pairs.

### 3.4 sRNARFTarget’s performance on IntaRNA 2.0’s testing data [31]

We took the confirmed interacting sRNA-mRNA pairs provided by [31]. Out of 160 confirmed interacting pairs, we excluded those pairs present in our training data and used the remaining 119 interacting pairs (88 pairs of *E. coli* (NC 000913) together with 31 pairs of *Salmonella* (NC 003197)) to compare the performance of sRNARFTarget with that or IntaRNA. We ran sRNARFTarget and IntaRNA for 17 sRNAs and 4240 mRNAs of *E. coli* and, 7 sRNAs and 4450 mRNAs of *Salmonella*. The final number of pairs was 102,385 (71,427 pairs of *E. coli* and 30,958 pairs of *Salmonella*) for both programs.

Figure 10 shows the ROC curve of sRNARFTarget and IntaRNA. AUROC of sRNARFTarget is 0.61, and IntaRNA is 0.59. sRNARFTarget’s performance is comparable to that of IntaRNA in terms of AUROC. We plotted the ROC curves separately for *E. coli* and *Salmonella* for both programs to check the behaviour of the two bacteria independently. The performance for *E. coli* is comparable for both programs (AUROC 0.61 for sRNARFTarget and 0.62 for IntaRNA) (Supplementary Figure 12). For Salmonella, sRNARFTarget achieved an AUROC of 0.58 and IntaRNA achieved an AUROC of 0.51 (Supplementary Figure 13).

**Figure 10:**
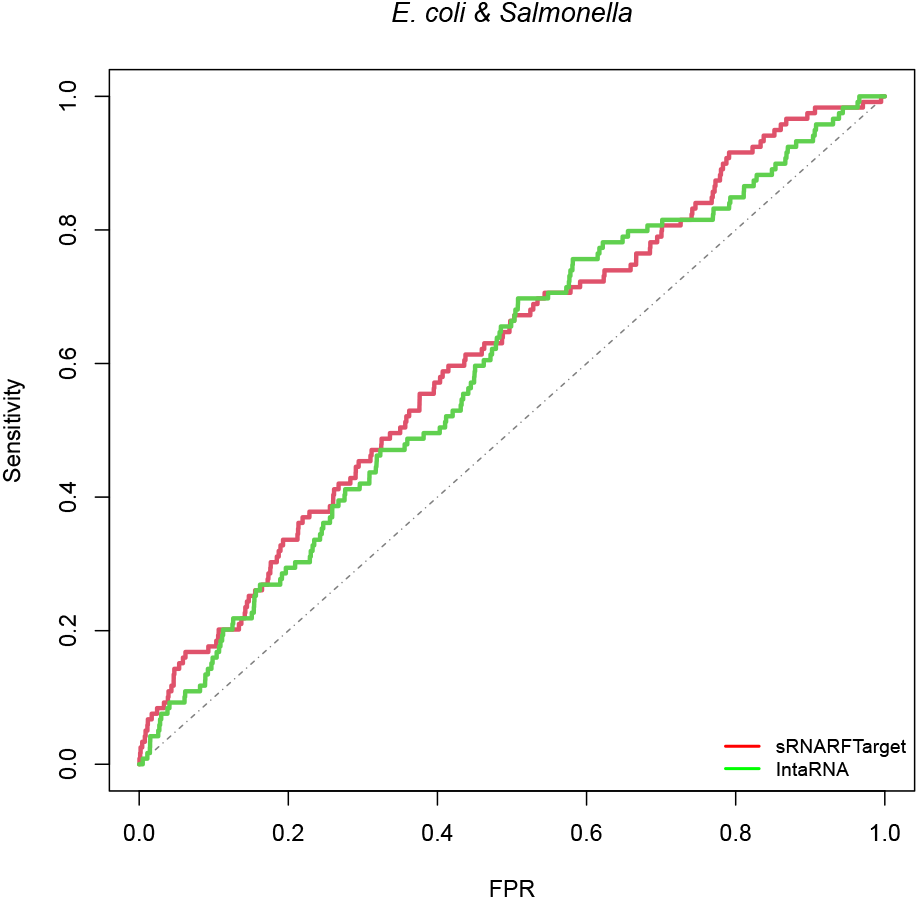
ROC curve for sRNARFTarget and IntaRNA on *E. coli* and *Salmonella* data. The plot shows the sensitivity (also called recall or true positive rate) as a function of the false positive rate (FPR). The dash line indicates random classifier performance.

Figure 11 shows the violin box plot for *E. coli* along with *Salmonella* for sRNARFTarget and IntaRNA. sRNARFTarget has a lower median rank compared to IntaRNA. P-value (Mann-Whitney test) indicates that the median rank of confirmed interacting pairs in sRNARFTarget is significantly lower than the median rank of IntaRNA.

**Figure 11:**
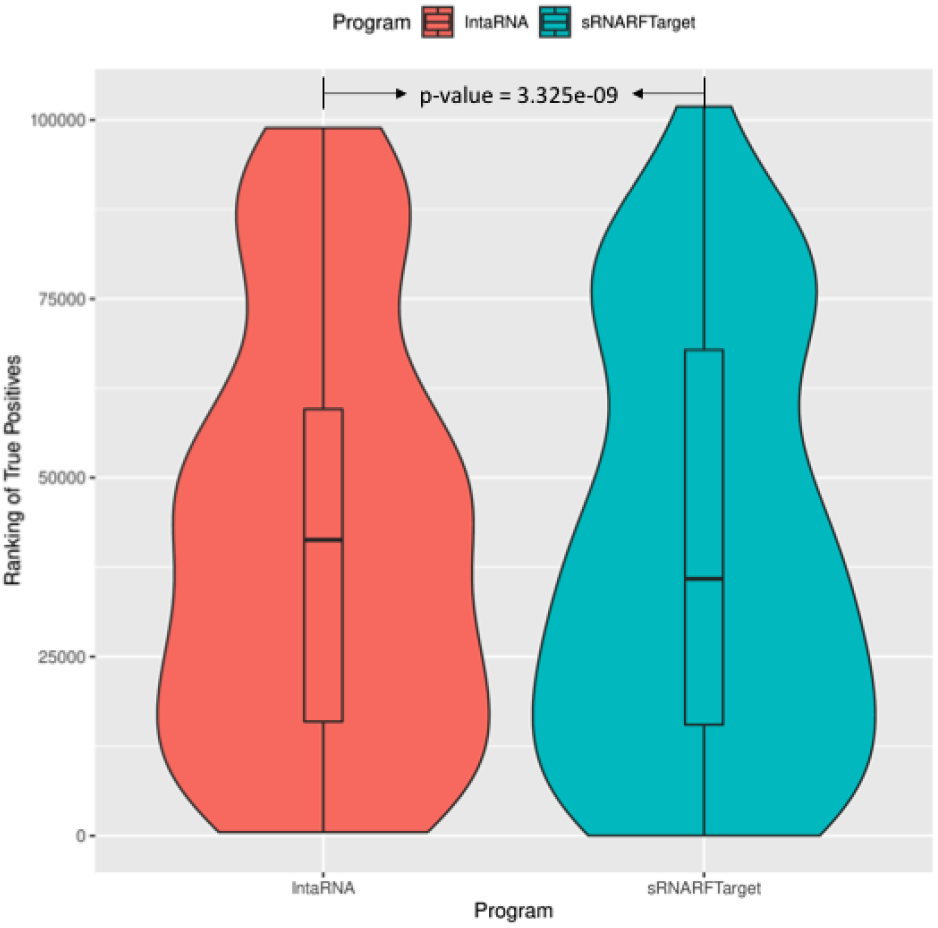
Rank (lower = better) distribution of 119 *E. coli* and *Salmonella* confirmed interacting pairs. The violin plot for each program shows the data density for different rank values and the horizontal line inside each box indicates the median rank of confirmed interacting pairs.

### 3.5 Programs execution time

In terms of execution time, sRNARFTarget is faster than CopraRNA and IntaRNA (Tables 6 and 7). Table 7 shows the time taken by the CopraRNA web server for job completion on selected sRNAs (CopraRNA is run for one sRNA at a time). These times were calculated by taking the difference between the job submission time and the job completion time (timestamp of job completion email). These times are not directly comparable to those shown in Table 6 as CopraRNA was run from the web server, and sRNARFTarget and IntaRNA were run from the Linux command line. sRNARFTarget execution time includes feature extraction (i.e., calculation of the trinucleotide frequency difference). To obtain interacting predictions for 1804 sRNA-mRNA pairs of *P. multocida*,sRNARFTarget took 31.4 seconds while IntaRNA took 6,196 seconds. To obtain interacting predictions for 93,280 sRNA-mRNA pairs (22 sRNAs and 4240 mRNAs) of *E. coli*, sRNARFTarget took 0.683% of the time taken by IntaRNA, which represents a 146-fold reduction in execution time (from more than 38 hours to 15 minutes). On average, sRNARFTarget is 100 times faster than IntaRNA with same or higher AUROC.

**Table 6:** Execution time for sRNARFTarget and IntaRNA on benchmarking data. Both programs were run on an Intel Core i7 (2.2 GHz) with 4 cores and 16 GB of RAM computer.

| Bacterium | No. of sRNAs/<br>mRNAs | Execution time<br>(HH:MM:SS) |  |
| --- | --- | --- | --- |
|  |  | sRNARFTarget | IntaRNA |
| *P. multocida* | 1/1804 | 0:00:31 | 1:43:16 |
| *Synechocystis* | 2/3179 | 0:01:18 | 2:33:02 |
| *Salmonella* | 7/4450 | 0:05:47 | 6:18:16 |
| *E. coli* | 22/4240 | 0:15:56 | 38:52:43 |
| **Average** | 8/3418 | 0:05:53 | 12:21:49 |

**Table 7:**
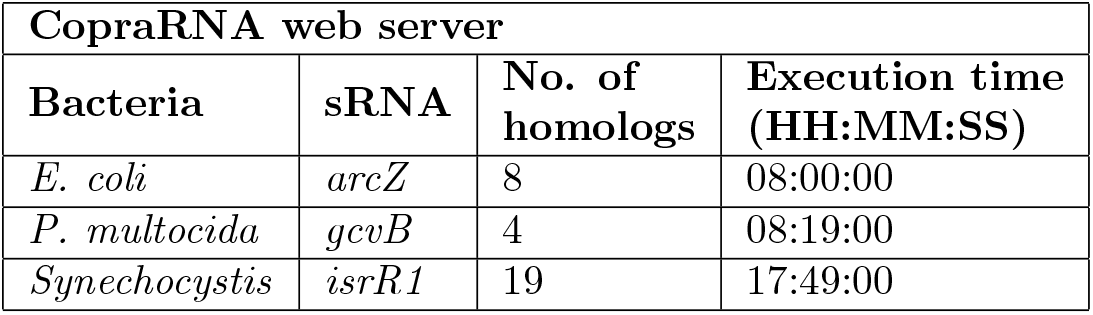
CopraRNA web server job execution time on selected sRNA for each bacterium on the benchmark data.

| CoprRNA web server |  |  |  |
| --- | --- | --- | --- |
| Bacteria | sRNA | No. of homologs | Execution time<br>(HH:MM:SS) |
| *E. coli* | *arcZ* | 8 | 08:00:00 |
| *P. multocida* | *gcvB* | 4 | 08:19:00 |
| *Synechocystis* | *isrR1* | 19 | 17:49:00 |

## 4 CONCLUSION

In this study, we present a transcriptome-wide sRNA target prediction program, sRNARFTarget. We collected experimentally verified sRNA-mRNA pairs from the literature to create a training data set consisting of 745 confirmed interacting sRNA-mRNA pairs. As a comparison, RNAInter [26] contains 408 sRNA-mRNA interactions. We selected a Random Forest model as the final model for sRNARFTarget using the trinucleotide frequency difference between sRNA-mRNA as features.

In our benchmark, we compared sRNARFTarget with CopraRNA and IntaRNA. Our results show that the comparative genomics-based approach used by CopraRNA is the best performing approach in terms of AUROC. However, unlike CopraRNA, sRNARF-Target does not require an sRNA or mRNA sequence to be conserved among other bacteria and can generate predictions for any number of sRNA and mRNA sequences. We also show that sRNARFTarget (the first ML-based approach used for this task) is 100 times faster (Table 6) than the best non-comparative genomics program available, IntaRNA, with better accuracy (Table 5). Another advantage of sRNATarget is its simplicity of use, as sRNARFTarget does not require any parameter setting; while IntaRNA has about a dozen parameters that need to be set [42]. As CopraRNA is the most accurate of the three programs, we suggest using CopraRNA when the homologs of the sRNA-mRNA sequences are available in at least four bacterial species. For transcriptome-wide prediction or when homolog sequences are not available, we recommend using sRNARFTarget.

## 5 ACKNOWLEDGEMENTS

This research was partially supported by a grant from the Natural Sciences and Engineering Research Council of Canada (NSERC) to L.P.-C. (Grant number RGPIN: 2019-05247). K.N. was partially supported by funding from Memorial University School of Graduate Studies. This research was enabled in part by support provided by ACENET (www.ace-net.ca/) and Compute Canada (www.computecanada.ca).

