## Supplemental Figures for "sRNARFTarget: A fast machine-learning-based approach for transcriptome-wide sRNA Target Prediction"

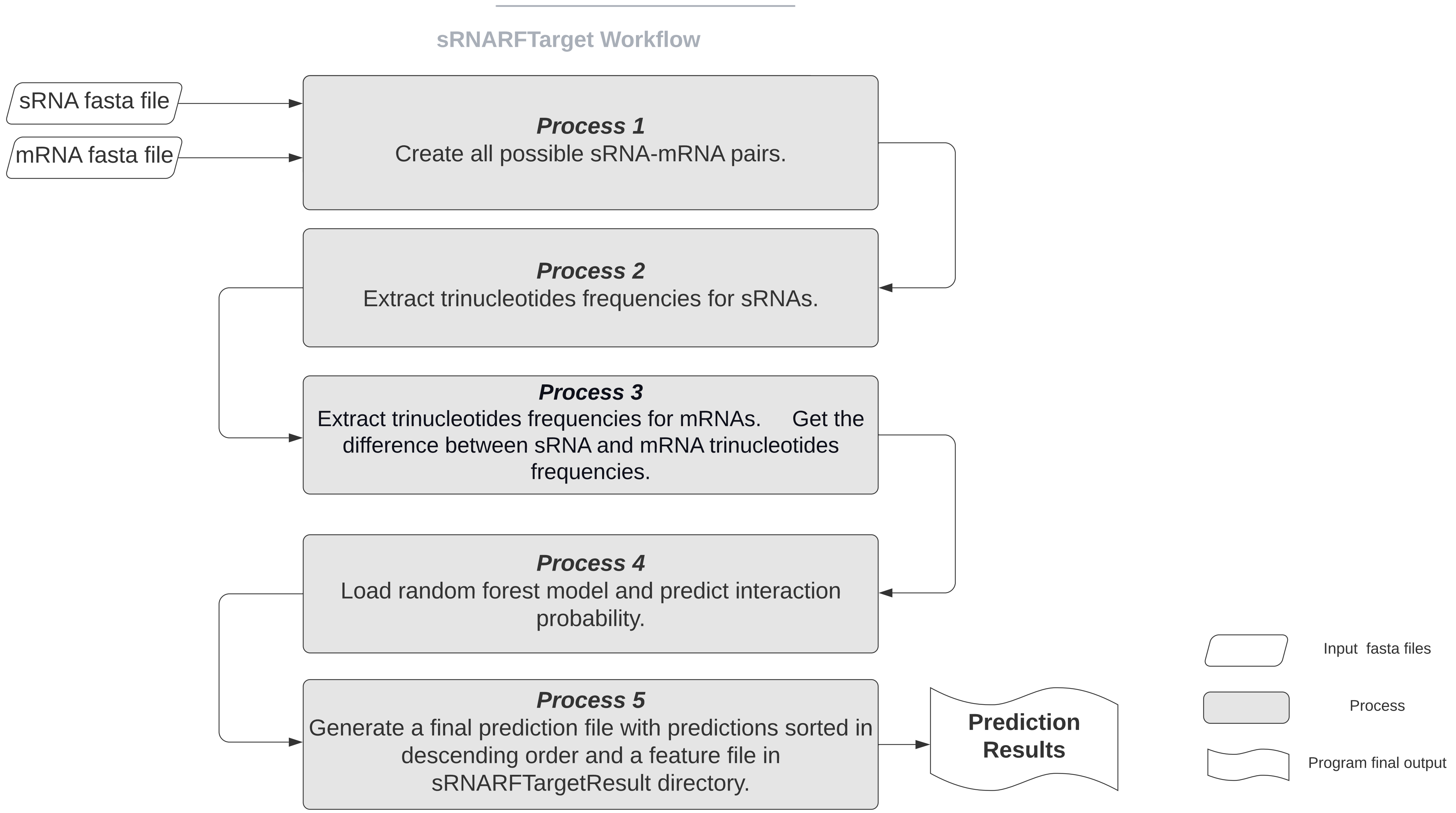


Supplementary Fig. 1

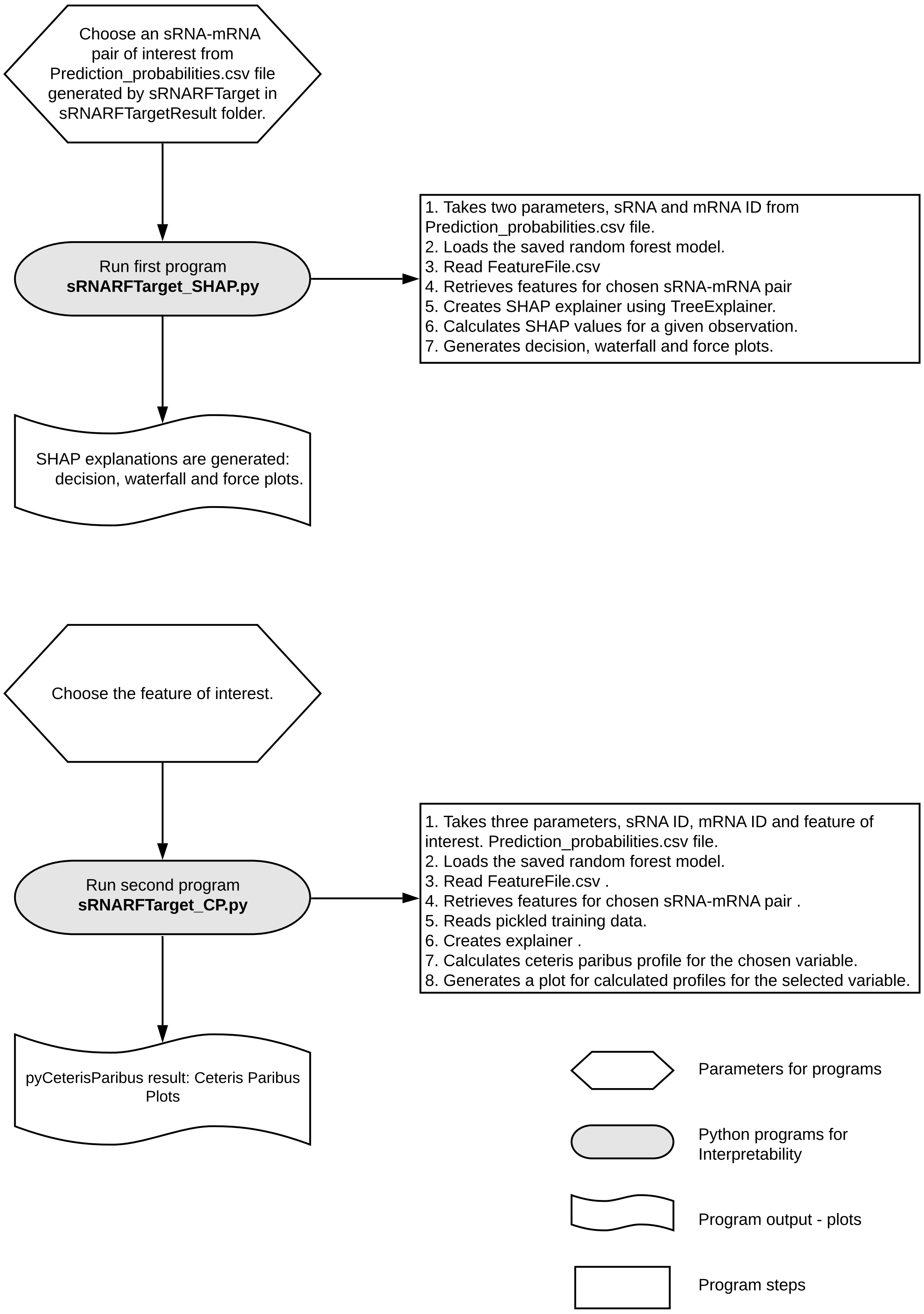


Supplementary Figure 2.

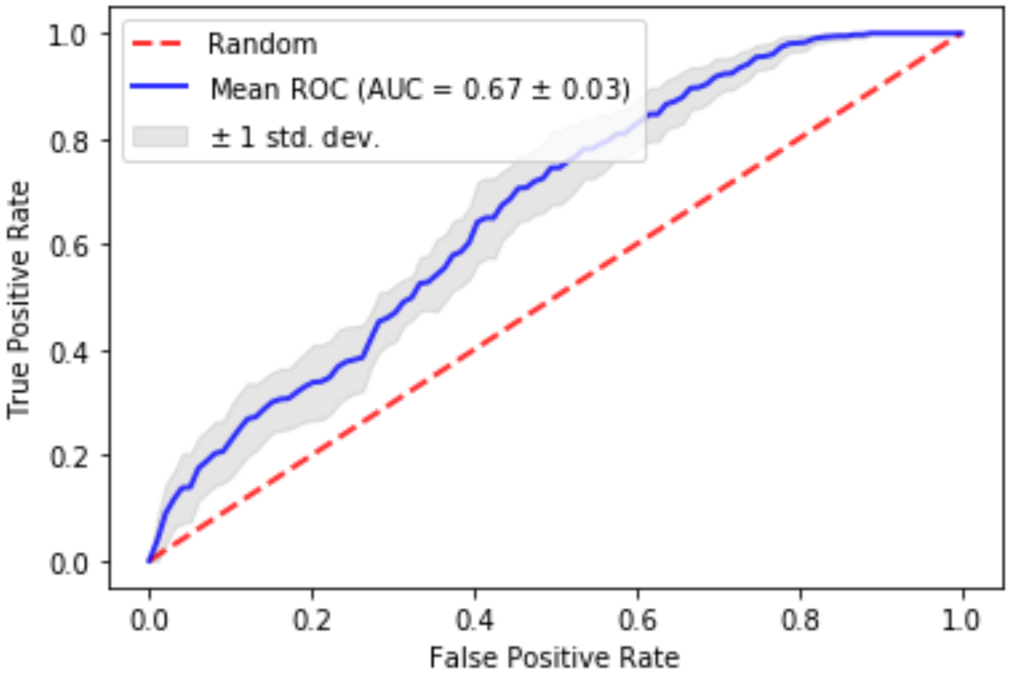


```
mean Average Precision=0.65
std Average Precision=0.04
```

Supplementary Fig. 3

Supplementary Fig. 4

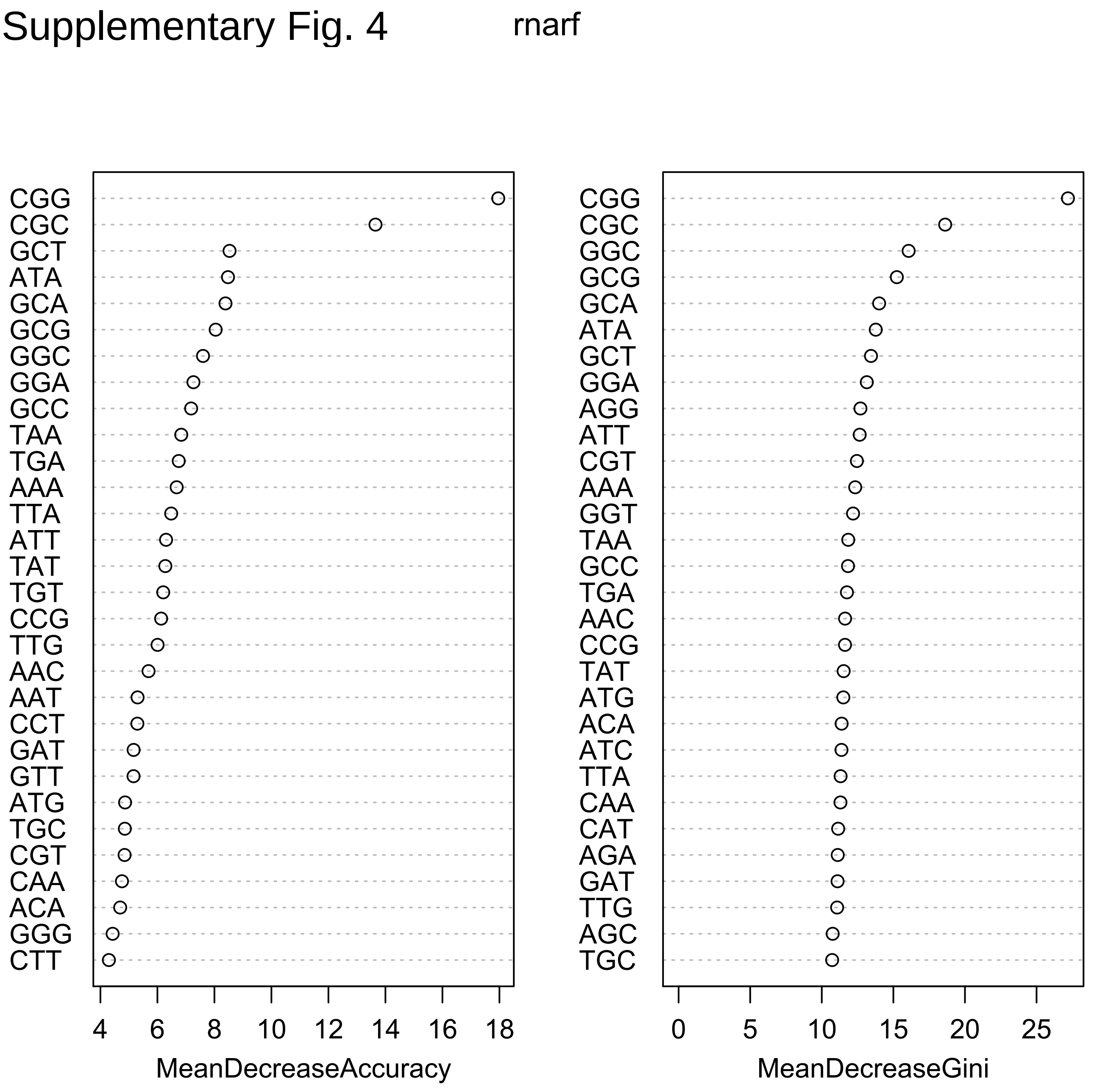

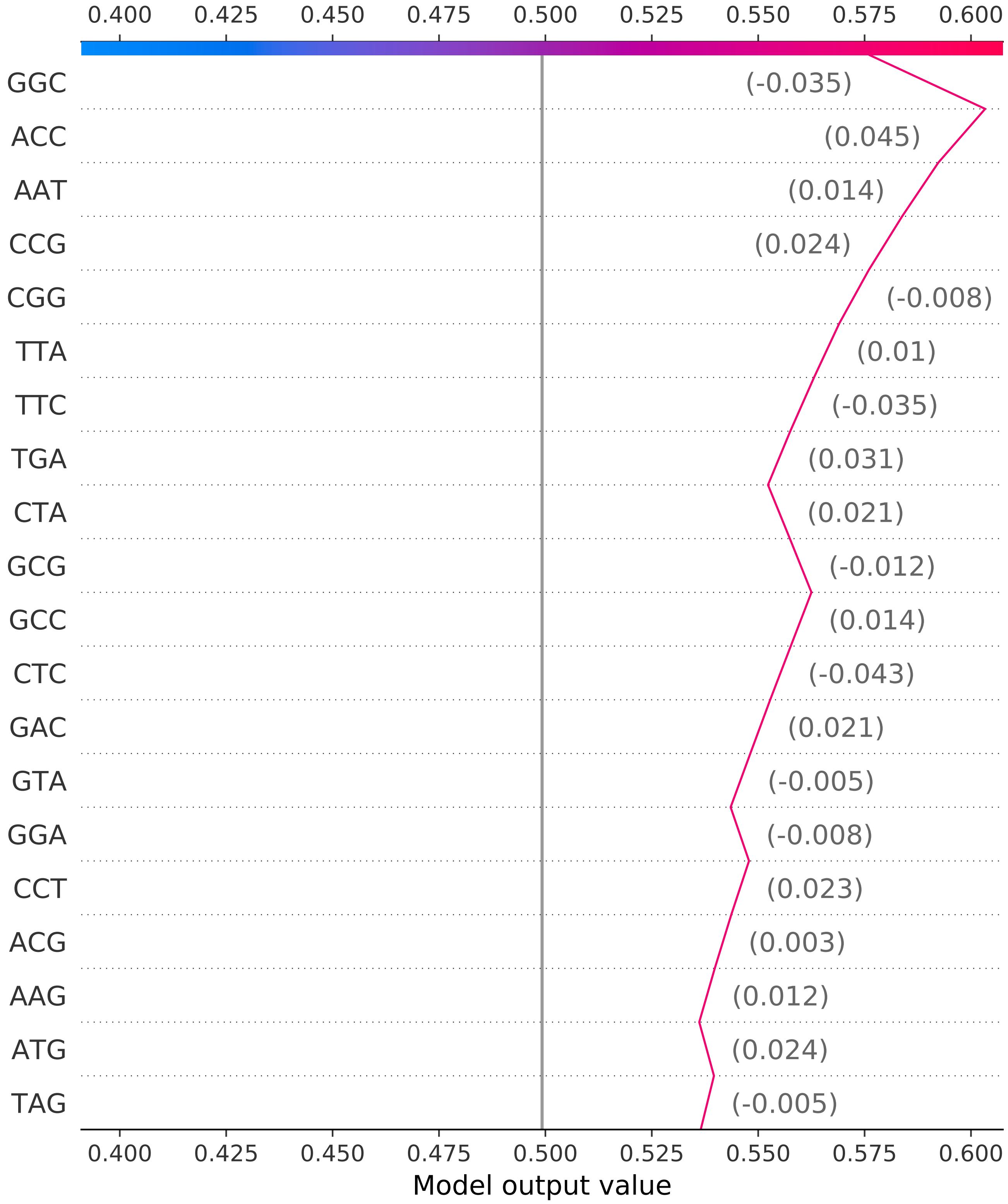


Supplementary Fig. 5

Supplementary Fig. 6

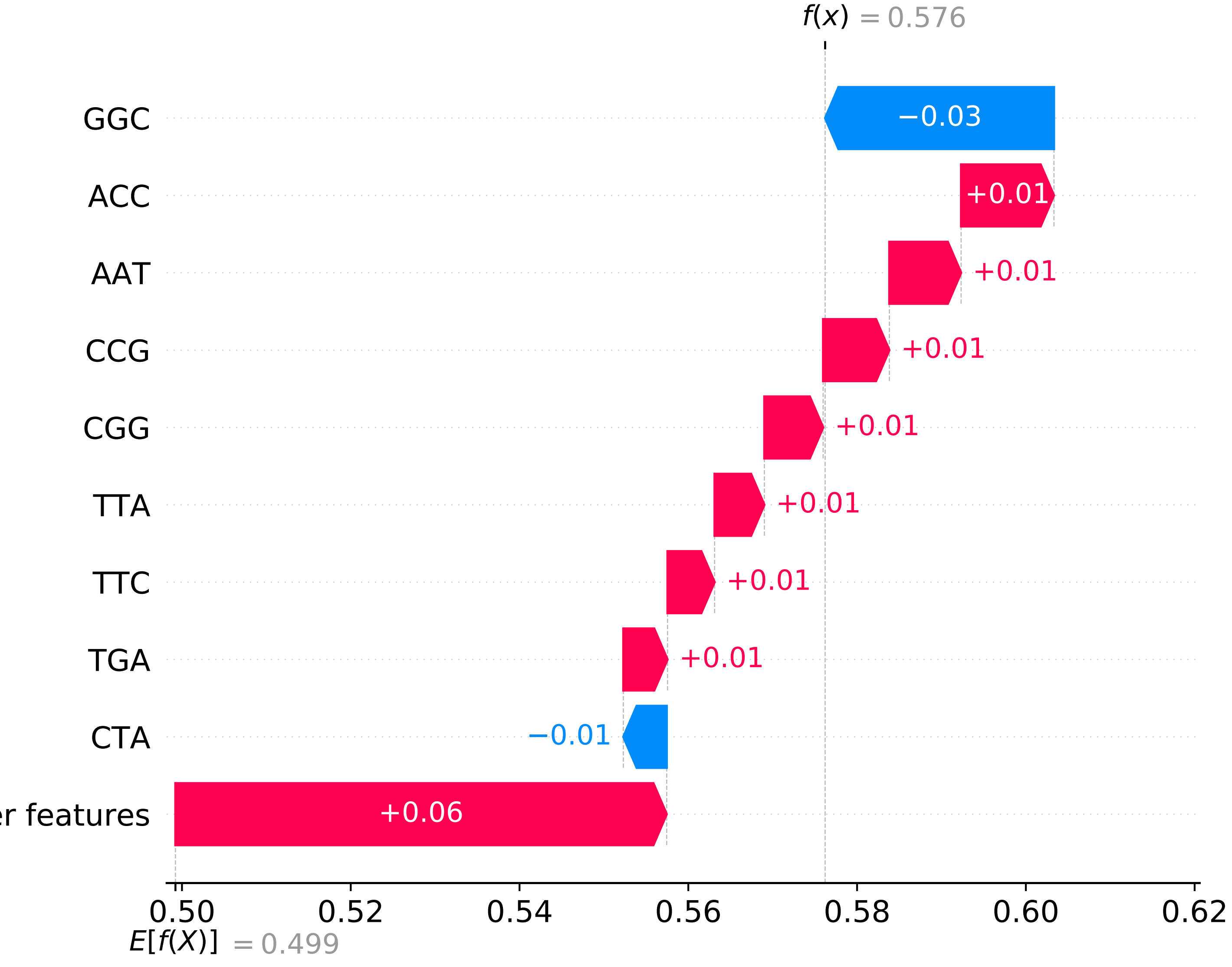

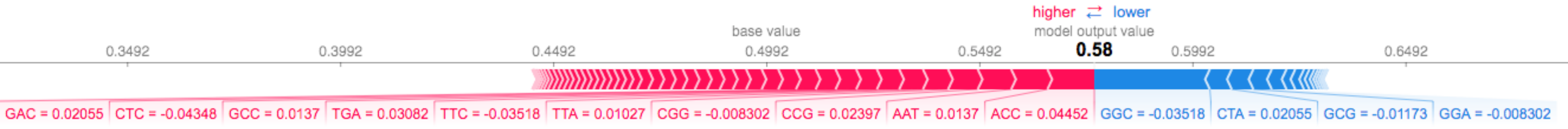


Supplementary Fig. 7

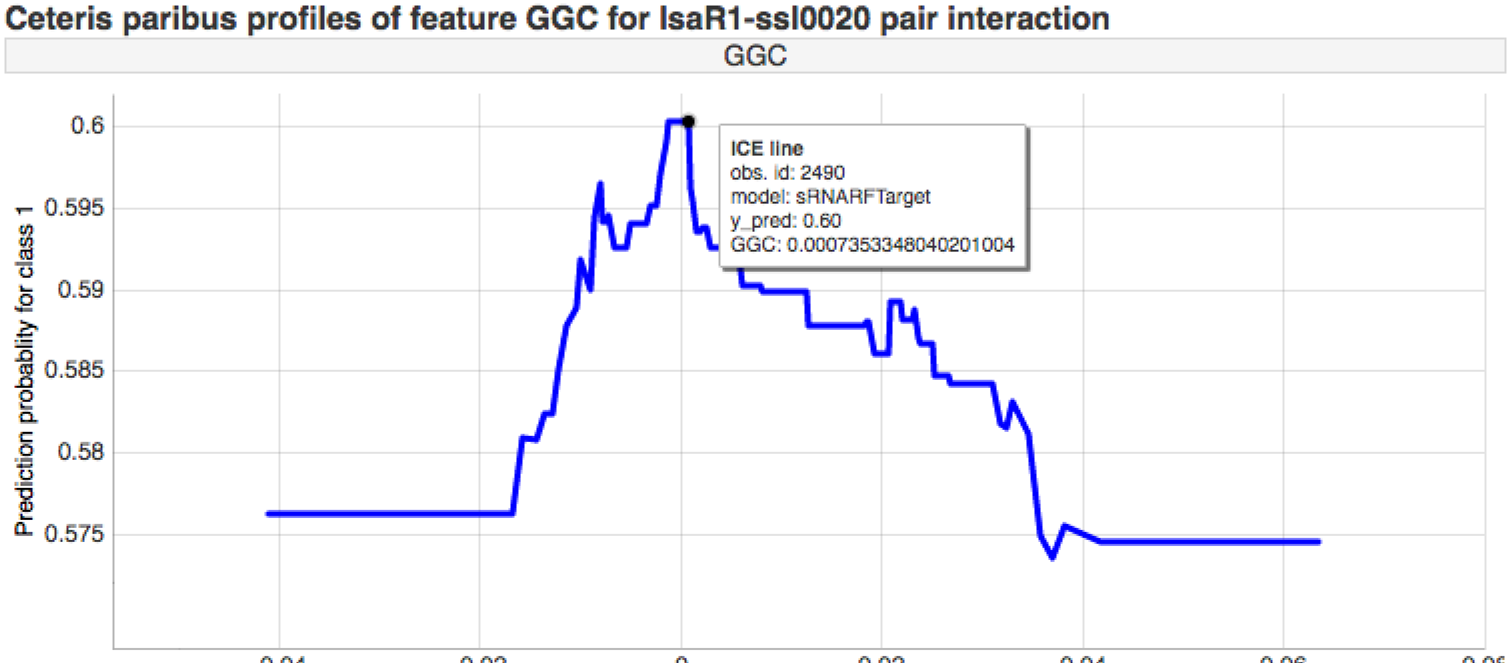


Supplementary Fig. 8

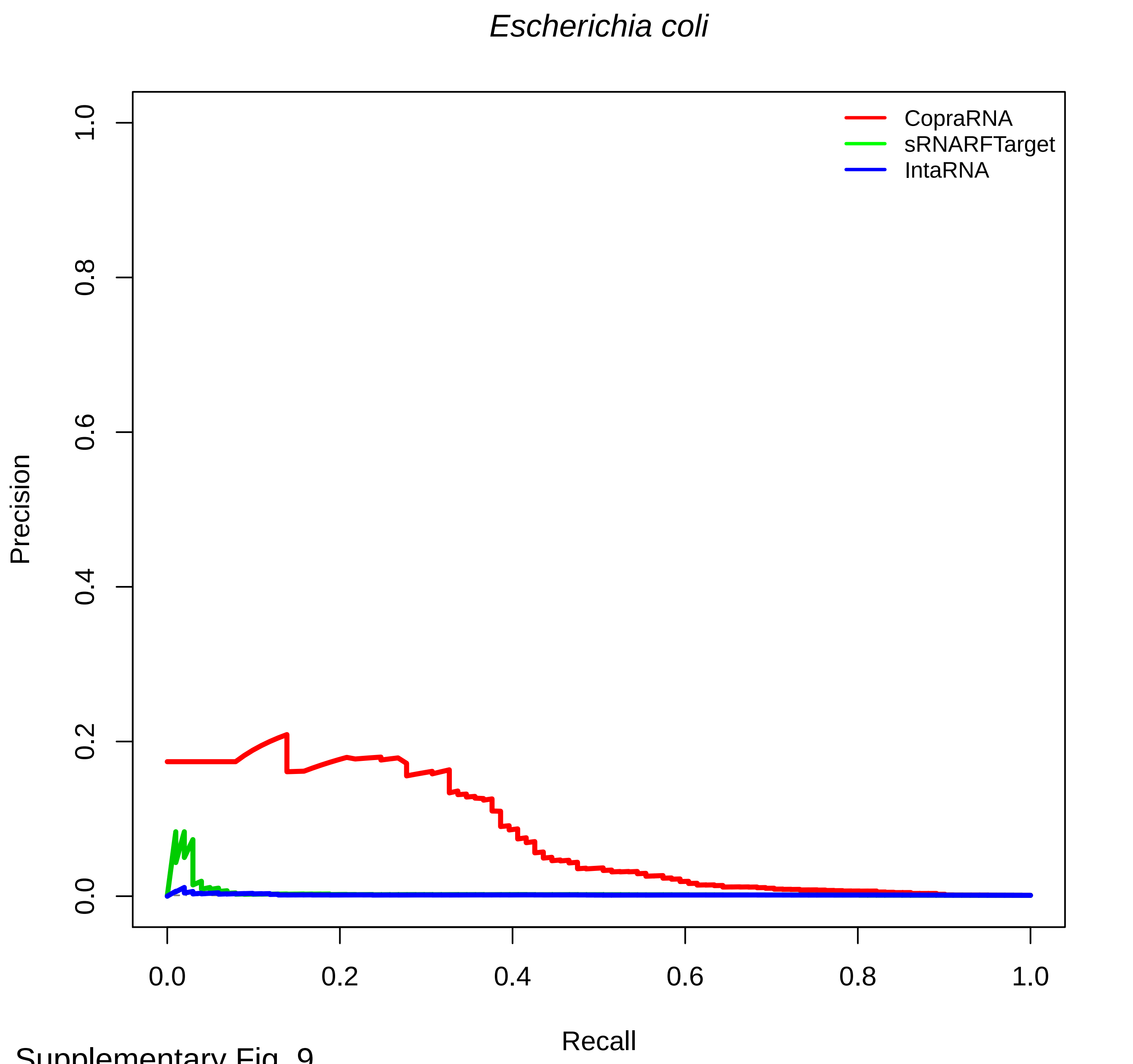


Supplementary Fig. 9

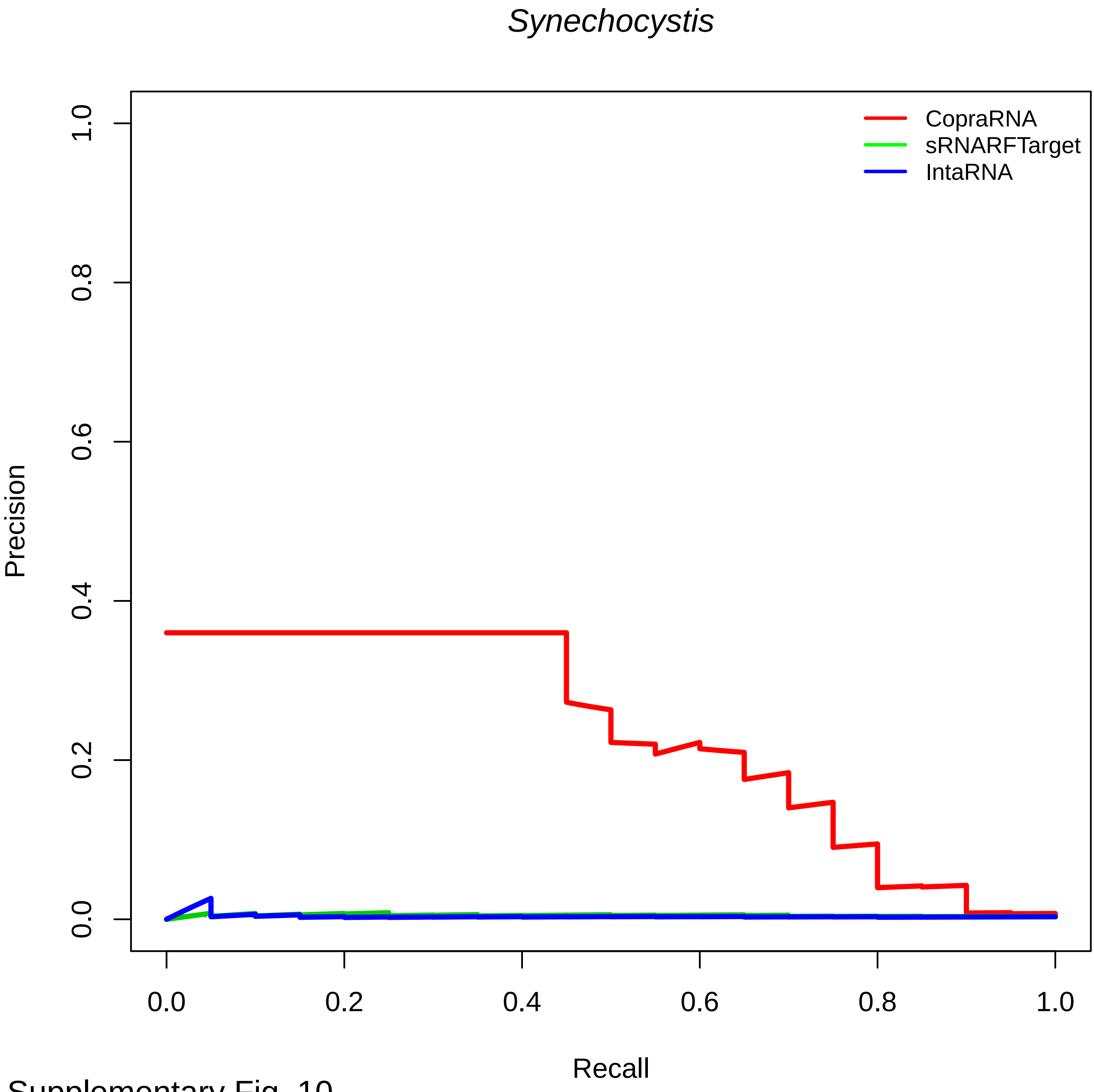


Supplementary Fig. 10

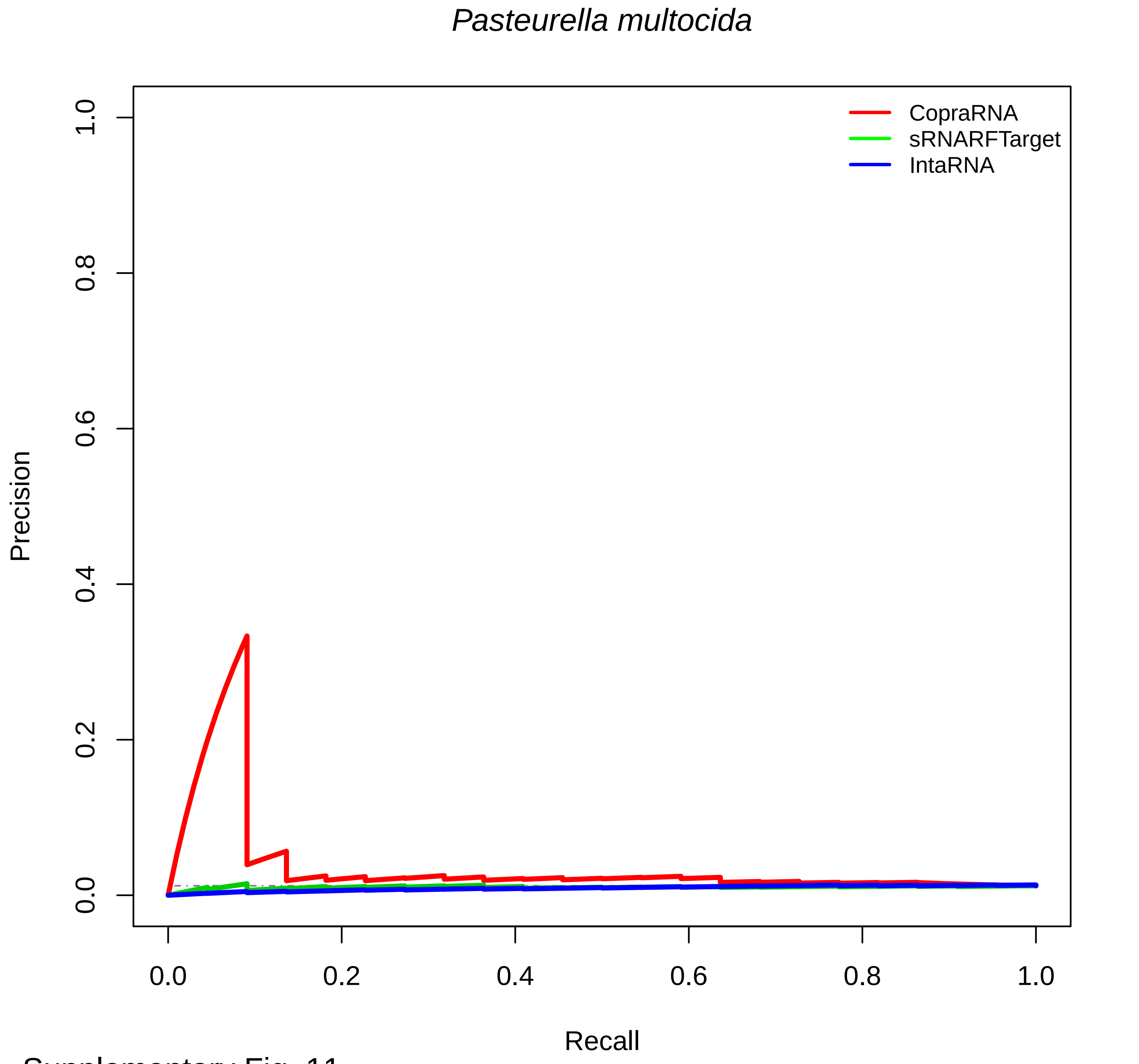


Supplementary Fig. 11

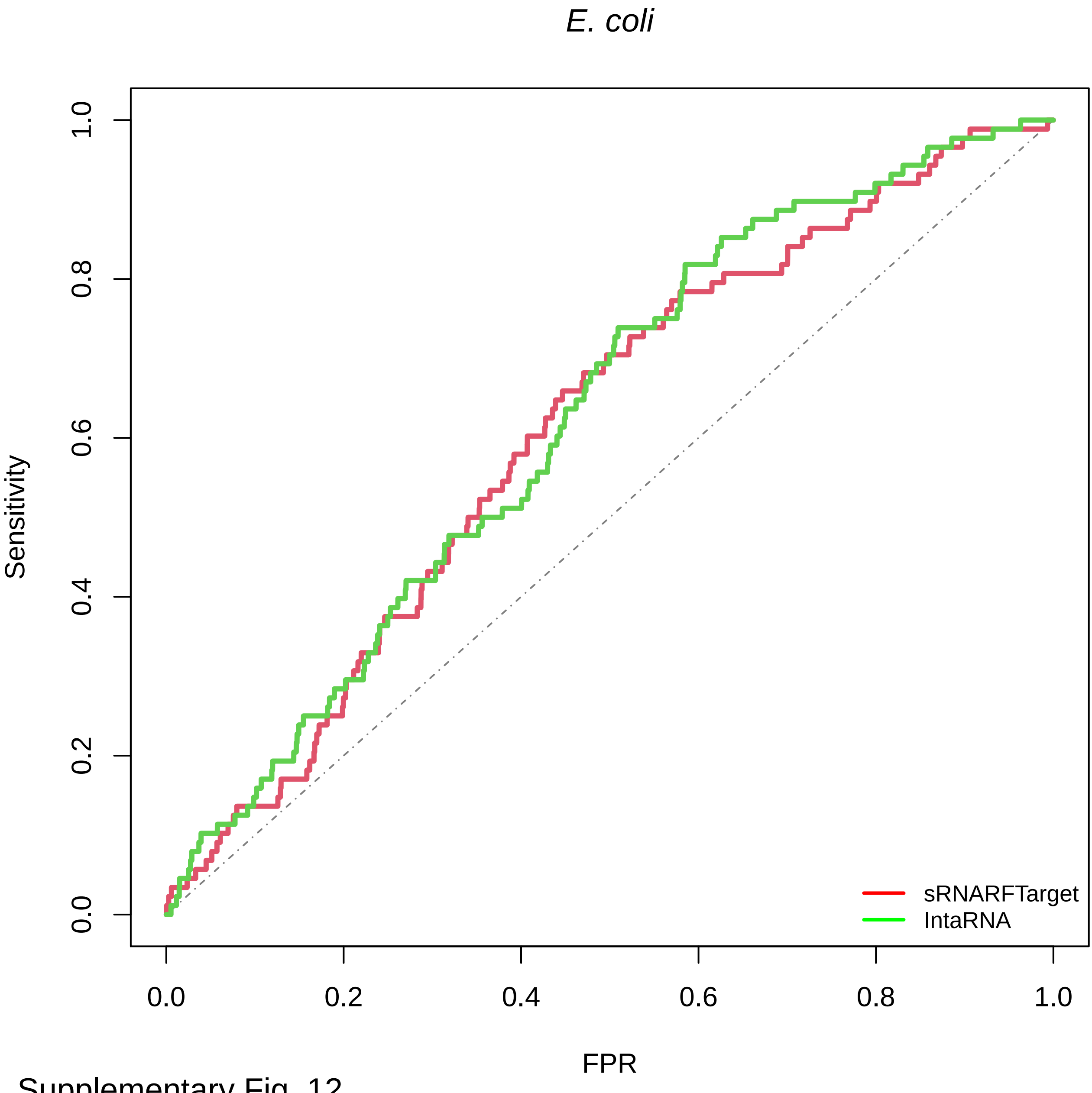


Supplementary Fig. 12

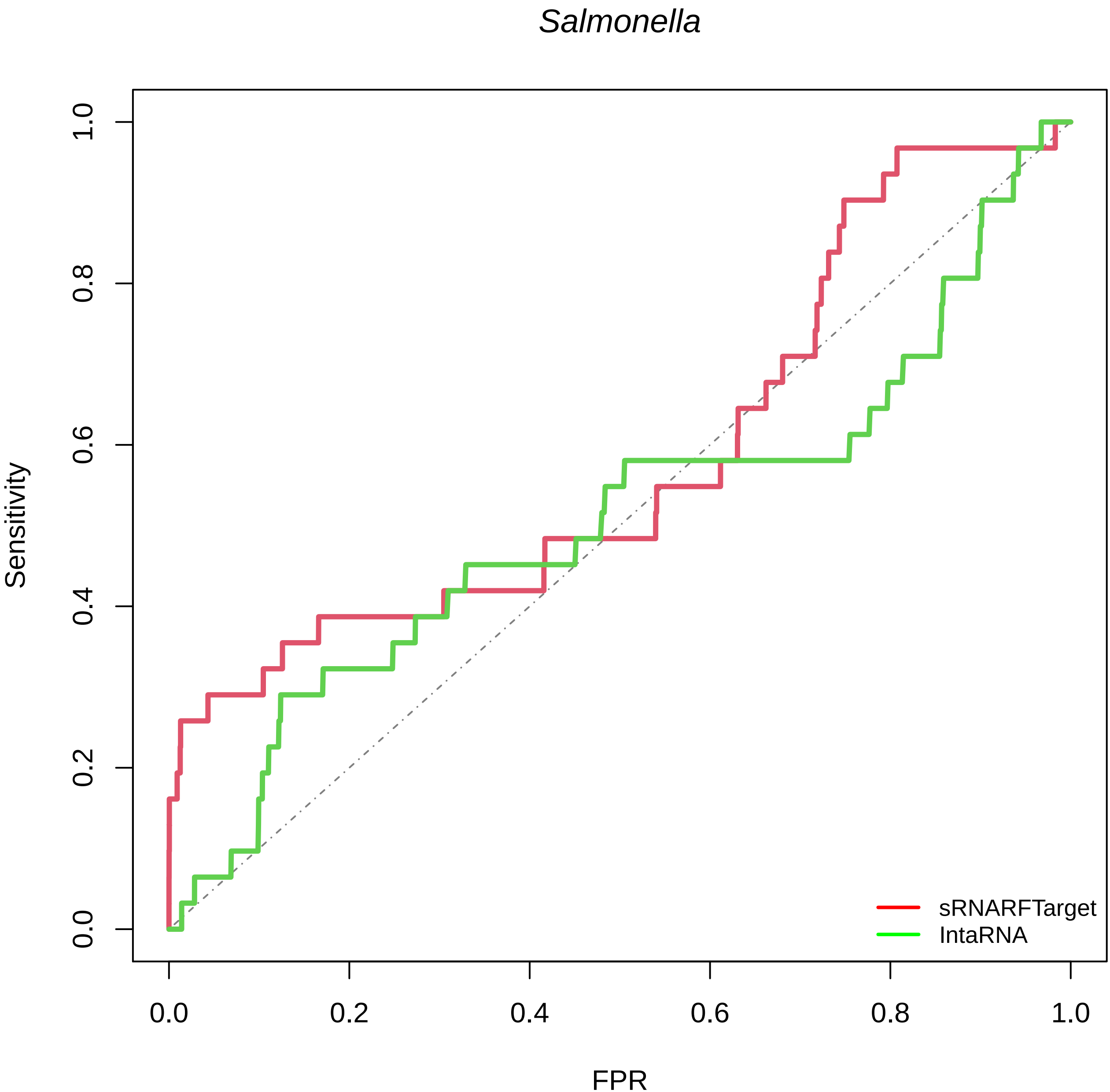


Supplementary Fig. 13
